# Comparative transcriptomics analysis of histone deacetylases, transcription factors, and ion channel genes in human iPSC-cardiomyocytes vs. the adult human heart

**DOI:** 10.64898/2026.01.16.700033

**Authors:** Maria R. Pozo, Michael P. Pressler, Anelia Horvath, Emilia Entcheva

## Abstract

Epigenetic modulators such as histone deacetylases (HDACs) and histone acetyltransferases (HATs) are known master regulators of gene expression that substantially impact cardiac electrophysiology. Novel pharmacological agents, HDAC inhibitors, are rapidly emerging as treatments for cancer and immune diseases, and their effects on cardiac ion channels (ICs) are of great interest. We used small interfering RNAs to individually suppress each of the known HDACs, including sirtuins (SIRTs), in human induced pluripotent stem-cell-derived cardiomyocytes (hiPSC-CMs), iCell2. Follow-up deep-sequencing allowed comparison to identically processed and normalized RNA sequencing data from adult human left ventricle (LV) from the GTEx database. The transcriptomics analysis revealed high similarity of gene expression patterns for cardiac ICs (with some differences in calcium influx and calcium buffering related genes), as well as strong co-regulation by cardiac transcription factors (TFs) and *HDACs/SIRTs* in both hiPSC-CMs and the adult LV. Partial least square regression models helped visualize links between HDACs/HATs, TFs, and cardiac ICs and helped identify potential key regulators of cardiac IC transcription. Powerful TFs, including *MEF2A, GATA4, 6* exerted positive effect on IC genes while *RUNX1* and *SHMT2* were distinct negative regulators in both sample types; *TRIM28* was found to serve opposite roles in the two sample types. In functional measurements, *HDAC* suppression primarily increased excitability, while *SIRT* suppression decreased excitability, in line with transcriptomic links. Our analysis offers insights about the role of epigenetic modifiers in regulating cardiac electrophysiology and informs the utility of hiPSC-CM as a scalable experimental model for cardiotoxicity testing of HDAC inhibitors.

## INTRODUCTION

The recent drive for new approach methods (NAMs) in scientific research seeks to improve human-specific tools for biomedical studies (1, 2). In cardiac work, this is especially relevant as there is a need for species-specific preclinical models that can faithfully capture electrophysiological behavior and drug responses in humans.

One approach for such studies is to utilize *in vitro* models such as human induced pluripotent stem-cell-derived cardiomyocytes (hiPSC-CMs). Human iPSC-CMs are a powerful tool for cardiac research and have been used extensively in probing cardiac electromechanics and metabolism (3–7). Their scalability makes them distinctly advantageous for high-throughput drug screening and, because they are patient derived, they are also well-suited for applications in personalized medicine. As an experimental cardiac model they are amenable to precise gene modulation techniques, including CRISPR-based approaches for human functional genomics studies (8–12). They can be combined with high-throughput optical approaches for comprehensive assessment of cardiac function (5, 6, 9, 13, 14) and such data are important in development of computational models of human cardiac responses (15). All of these make hiPSC-CMs usage a promising opportunity for clinically relevant advancements (7, 16–18).

It is important to understand whether mechanistic studies conducted in human iPSC-CMs are translatable to *in vivo* scenarios and if similar molecular drivers and relationships are at play. In evaluating hiPSC-CM as a model of the adult human heart, recent work has shed light on whole-transcriptome level characterization of high performing, commercially available hiPSC-CM cell lines (19, 20) and whole-transcriptome variances between these cell lines and the adult heart (21, 22). However, in-depth analyses of gene expression relevant to cardiac electrophysiology, directly comparing the hiPSC-CM and the adult heart, have been lacking. One challenge is that healthy cardiac tissue samples are a scarce resource, making large sample-size studies difficult. To address this, we accessed a comprehensive public dataset of genomic and transcriptomic profiles from hundreds of adult humans, the Genotype-Tissue Expression (GTEx) Portal (23) and extracted left ventricle (LV) RNA sequencing data. To our knowledge, this report is the first to directly compare hiPSC-CM transcriptomics to the GTEx resource.

For this study, we were particularly interested whether hiPSC-CM are a suitable model to study the effects of epigenetic modulators. The complex epigenetic landscape of the heart is closely tied to cardiac electrophysiology and pathology (24–27). Regulation by classes of epigenetic modulators, namely histone deacetylases (HDACs) and histone acetyltransferases (HATs), plays a role in cardiac electrophysiology and mechanics, by canonical regulation of chromatin organization (28–31) and by non-canonical action on other proteins (32, 33), **Fig. 1A**. While the rapid development of pharmacological HDAC inhibitors (HDACi) to treat cancer and autoimmune diseases prompted a focus on screening for potential cardiotoxic effects (19), HDACis are also, paradoxically, being investigated as potential cardiac therapeutics, with evidence of cardioprotection against ischemia/reperfusion injury and other beneficial roles (34–36). The utility of human iPSC-CMs in the development of new HDACis was recently reviewed (37) and, importantly, the identification and development of human-centered NAMs for cardiac epigenetics research is critical to circumvent issues with interspecies epigenomic variation (38, 39) and cell and species-specific ion channel response variation (40, 41).

**Figure 1.**
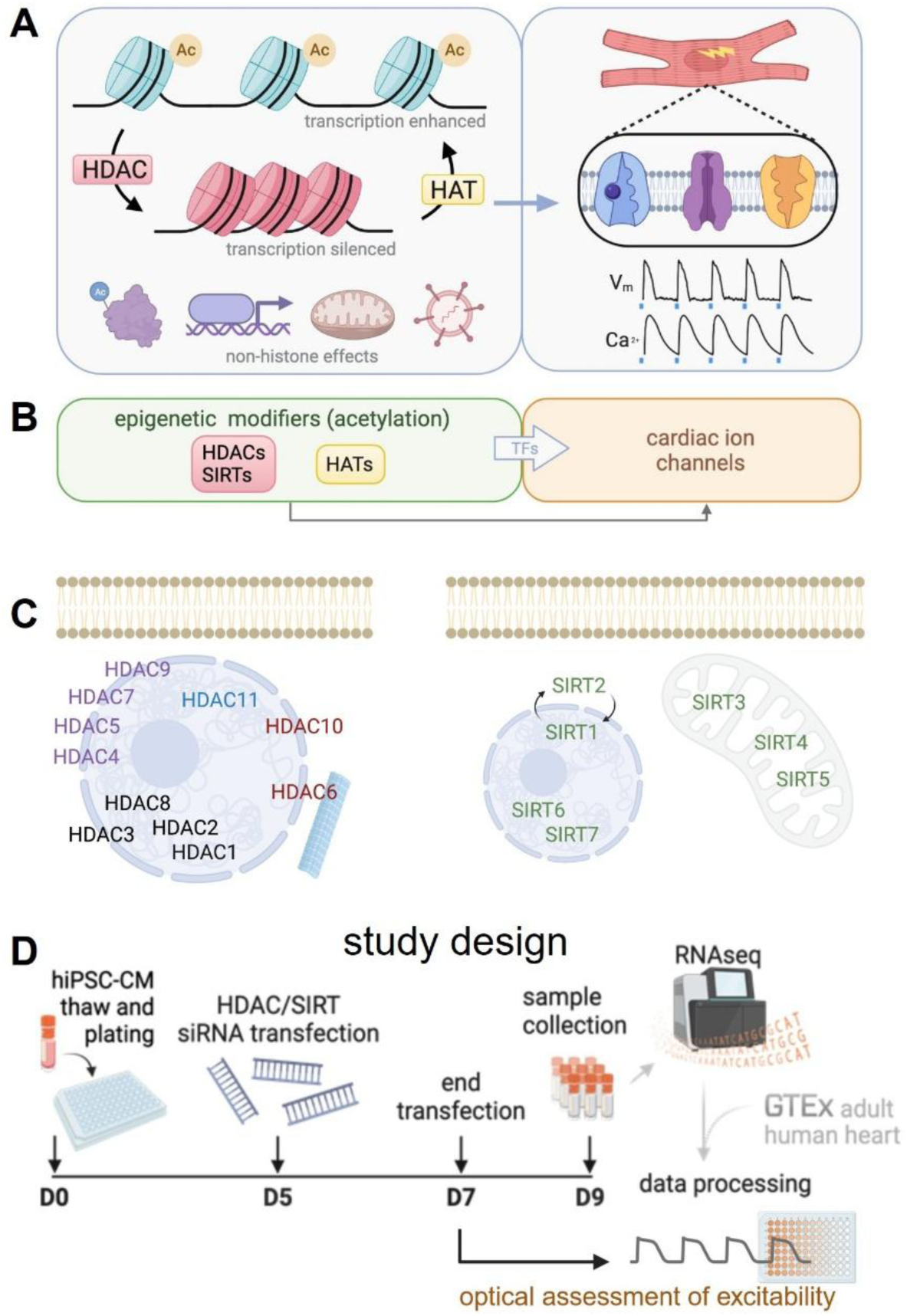
Epigenetic de-acetylation modifiers and cardiac ion channels: experimental design to compare their role in human iPSC-CMs and adult human cardiac tissue. **A**. HATs and HDACs epigenetically regulate gene expression through reversible (de)acetylation of histone proteins and through direct action on TFs with potential regulatory effects on cardiomyocyte electrophysiology. In additional to catalytic action on chromatin compaction/relaxation, the HDACs and HATs have also non-canonical roles as protein stabilizers, chaperons, metabolic modulators, regulators of viral entry. **B.** These relationships were modelled computationally with HDAC/SIRT and HAT expression as inputs, cardiac ion channel expression as outputs, and TF expression as intermediaries. **C.** Cellular localization of HDACs and SIRTs corresponds to roles in chromatin remodeling, protein interaction, mitochondrial regulation, etc. Different classes of HDACs and SIRTs are color-coded. **D.** Experimental design and timeline for siRNA treatment includes RNA sequencing of hiPSC-CMs, which was processed alongside RNAseq data (adult human heart) accessed from the GTEx database. A separate set of 48 h siRNA treated live hiPSC-CM samples were used in assays for excitability.

Our group previously explored the GTEx LV dataset and reported on sex-dependent transcriptomic links between *HDAC/SIRTs* and cardiac electrophysiology in the human heart (42). In this study we used siRNA suppression of *HDAC/SIRTs* in hiPSC-CM samples and transcriptomic analysis of relevant genes, dissecting epigenetic relationships and evaluating their similarity to relationships found in the adult human heart.

## RESULTS

### Patterns of gene expression in hiPSC-CM vs. the adult human LV

To better understand the transcriptome of electrophysiology-related genes in hiPSC-CM, especially in comparison to the adult LV (obtained from the GTEx database), RNAseq data from both sample types were transformed from simplex/Aitchison space into Euclidian space, allowing direct quantitative comparison of gene expression profiles (as described in Methods and in our previous study (42). Relative expression levels of key cardiac ion channel genes were obtained, along with relative expression levels of a larger selection of important electrophysiology genes, termed the cardiac “rhythmonome” (43). The overall pattern of baseline expression of key cardiac ion channels was remarkably similar between hiPSC-CM and adult LV, **Fig. 2A**. This finding was pervasive when comparing the rhythmonome genes (**Fig. 2B**), HDAC-coding genes, cardiac HAT-coding genes, and cardiac TF genes, (**Fig. 3A-C**). The radial displays in **Figs. 2**-**3** help convey the degree of similarity, which is most obvious for the select cardiac ion channels, HATs, and cardiac TFs (difference scores of 0.157, 0.165, and 0.144, respectively).

**Figure 2.**
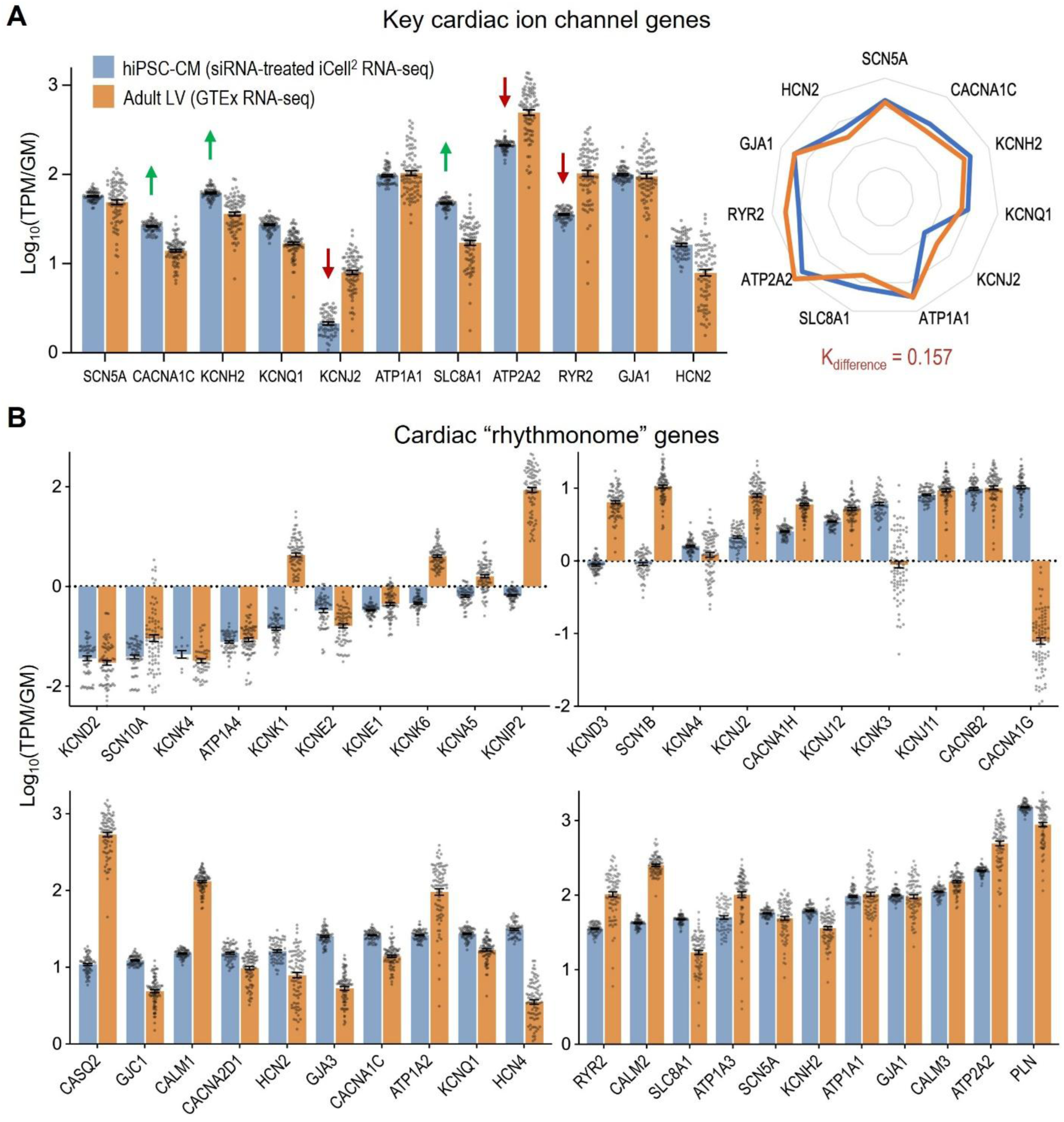
Comparative profiles of gene expression of cardiac ion channel genes in hiPSC-CMs and in human adult left ventricle. **(A)** Relative expression of select cardiac ion channel genes and **(B)** more comprehensive cardiac electrophysiology “rhythmonome” genes in hiPSC-CM (n=61 female) and adult LV (n=84 female, from GTEx) bulk RNA-sequencing data. The inset in panel A is a radial plot of the expression averages for the two groups to highlight pattern similarity of expression. K_difference_ is listed as well. Individual data points displayed; error bars represent mean and SEM.

**Figure 3.**
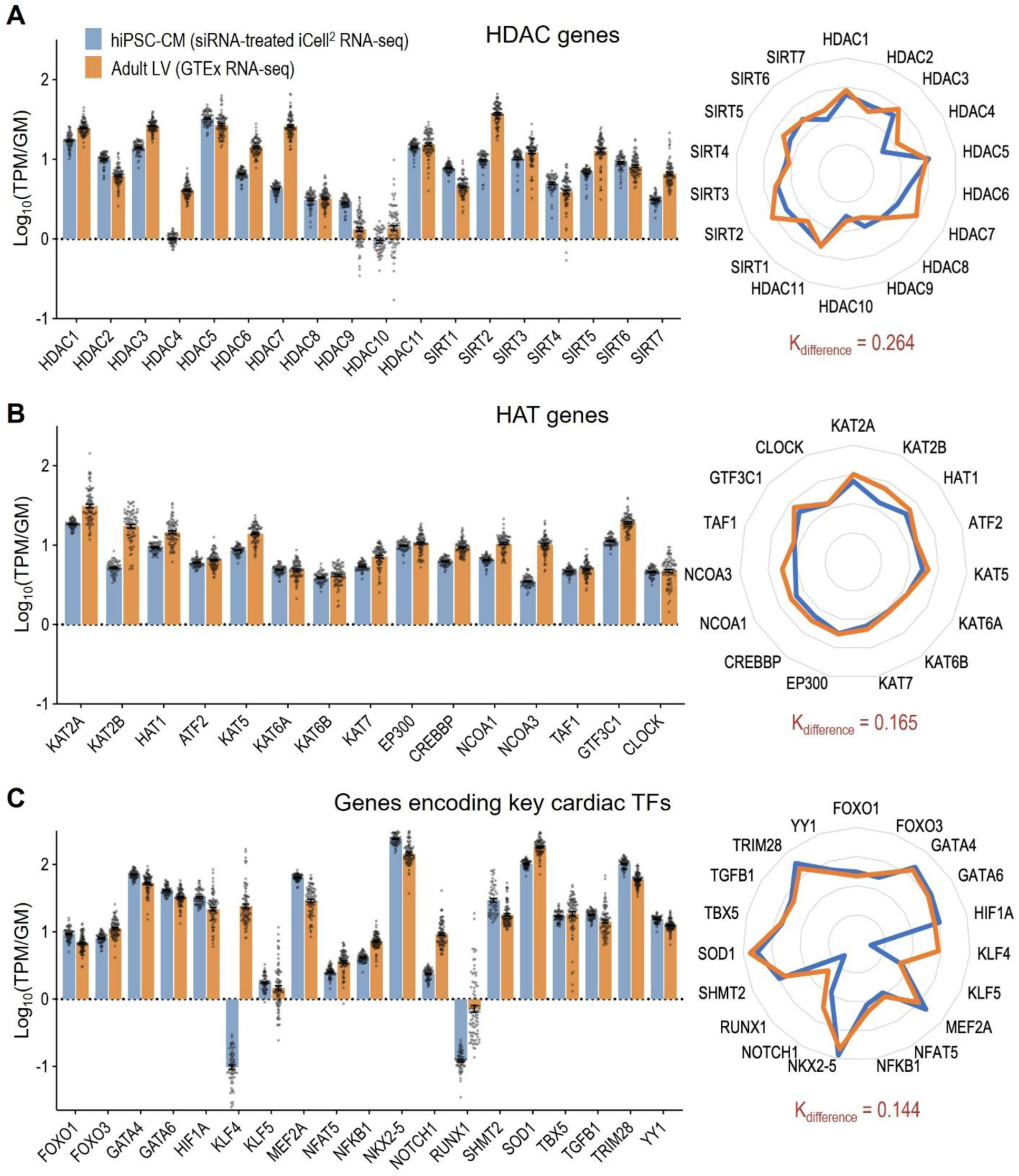
Comparative profiles of gene expression of HDACs/SIRTs, HATs, and TFs in hiPSC-CMs and in human adult left ventricle. Relative gene expression of *HDAC*s/*SIRT*s (**A**), *HAT*s (**B**), and key cardiac transcription factor genes (**C**) in hiPSC-CM (n=61) and adult LV (n=84, from GTEx) bulk RNA-sequencing data. The radial plots of the expression averages for the two groups are shown to the right of panels A, B and C to highlight pattern similarity or differences of expression. K_difference_ is listed for each gene set. Individual data points displayed; error bars represent mean and SEM.

Some interesting differences are captured here as well. Genes for ion channels providing calcium entry into the cell, like *CACNA1C* (L-type Ca^2+^), *SLC8A1* (NCX), and *CACNA1G* (T-type Ca^2+^) were relatively upregulated in hiPSC-CM compared to the adult LV (see radial plot inset of **Fig. 2A**). The same was true for repolarization related K^+^ channel genes *KCNH2* and *KCNQ1.* The genes underlying the pacemaking “funny” current I_f_, *HCN2* and *HCN4*, were both expressed at higher relative levels in the hiPSC-CMs, which may contribute to their automaticity (**Fig. 2B**)(44). The *HCN2:HCN4* ratio is known to increase with development and maturity (45). As expected, the adult human LV shows much higher *HCN2:HCN4* ratio compared to the hiPSC-CMs (**Fig. 2B**). While the main Cx43-encoding gene *GJA1* shows similar levels in hiPSC-CM and the adult LV, the hiPSC-CM have higher expression levels of *GJC1* and *GJA3* (coding for non-ventricular gap junctions Cx45 and Cx46), which normally localize to the conduction system in the heart.

Conversely, several ion channel genes were downregulated in hiPSC-CM compared to the adult LV. For *KCNJ2*, this is a well-known distinction that explains in part the more depolarized resting membrane potential in hiPSC-CMs compared to adult cardiomyocytes (46–49). Calcium handling genes *ATP2A2* (SERCA2 pump), and *RYR2*, *CALM1*, *CALM2*, *CASQ2* were also expressed at lower relative levels in the hiPSC-CMs compared to the adult human LV, with phospholamban (*PLN*) being higher expressed in hiPSC-CMs. This differential expression of *ATP2A2* and *PLN* could lead to lower contractility and inferior relaxation in hiPSC-CMs, consistent with less mature phenotype. Furthermore, lower expression of genes related to calcium buffering (calmodulins *CALM1, CALM2,* and calsequestrin *CASQ2*) combined with a less active SERCA pump and higher expression of calcium influx related genes may make hiPSC-CMs more prone to calcium overload. Other lower-expressed genes in the hiPSC-CMs included *KCNIP2*, an important contributor to the repolarizing transient outward current Ito, as well as *KCNK1* and *KCNK6* of the two-pore domain (2P) K^+^ channel family (TWIK-1 and TWIK-2, respectively). *KCNK3*, which codes for another 2P K^+^ ion channel (TASK-1) was significantly upregulated in the hiPSC-CMs compared to the adult LV.

From the *HDAC* and *SIRT* genes (**Fig. 3A**), with a difference score of 0.264, the most notable differential expression was seen in *HDAC4, HDAC7*, *SIRT2* and *SIRT7*, which are expressed at lower levels in the hiPSC-CMs than in the adult LV. Conversely, *SIRT1* shows higher relative expression in hiPSC-CMs, as does *HDAC9* (often considered the main cardiac HDAC (50)), although interestingly it is one of the lowest expressed HDACs in both the hiPSC-CM and adult LV.

Similar expression levels of histone acetyltransferase genes are seen in the hiPSC-CMs and in the adult LV, **Fig. 3B**. These act as key regulators of cell growth, stress response, rhythm, and metabolism through recruitment of TFs or as TFs themselves. Cardiac TFs follow very similar patterns of relative expression in the hiPSC-CMs and the adult LV (visualized in the radial plot, **Fig. 3C**). Several TF genes show higher expression in the hiPSC-CM compared to the adult LV, e.g. *MEF2A, NKX2.5, FOXO1* and *TRIM28*. The most striking discrepancy in the TF gene expression is seen in *KLF4*, which is one of the four Yamanaka factors in stem cell reprogramming, suppressed early on in cardiomyocyte differentiation but well expressed in the mature adult myocardium (51). For the main gene sets considered in this study, relative expression levels in cardiac muscle cells and cardiac ventricular fibroblasts were extracted from publicly available and preprocessed single-nuclei sequencing of human LV tissue (**Fig. S1**, (52)).

### Whole-transcriptome analysis upon HDAC/SIRT knockdown in hiPSC-CMs

While the human adult LV gene expression had natural variability due to a wide range of donors, we used siRNA perturbations (siRNAs listed in **Table S1**) of the *HDAC* and *SIRT* genes to uncover important relationships. Whole-transcriptome analysis of the *HDAC/SIRT* knockdowns in hiPSC-CM samples included our own UMAP clustering and differentially expressed gene (DEG) analysis performed by and provided by GENEWIZ/Azenta (South Plainfield, NJ). Knockdown of each *HDAC/SIRT* gene resulted in several discernable clusters (**Fig. 4A, B**). For example, knockdowns of *HDAC1*, *HDAC9*, *SIRT2*, and *SIRT3* formed distinct clusters, indicating that *HDAC1, HDAC9, SIRT2,* and *SIRT3* knockdown groups contain samples that are similar to each other at the level of the whole transcriptome and different from other groups. Some sample groups did not cluster well in the displayed projections, for example *HDAC11* knockdowns, which could be related to the UMAP parameters chosen or could be due to the dynamic nature of *HDACs* and *SIRTs* being difficult to capture in low dimensions.

**Figure 4.**
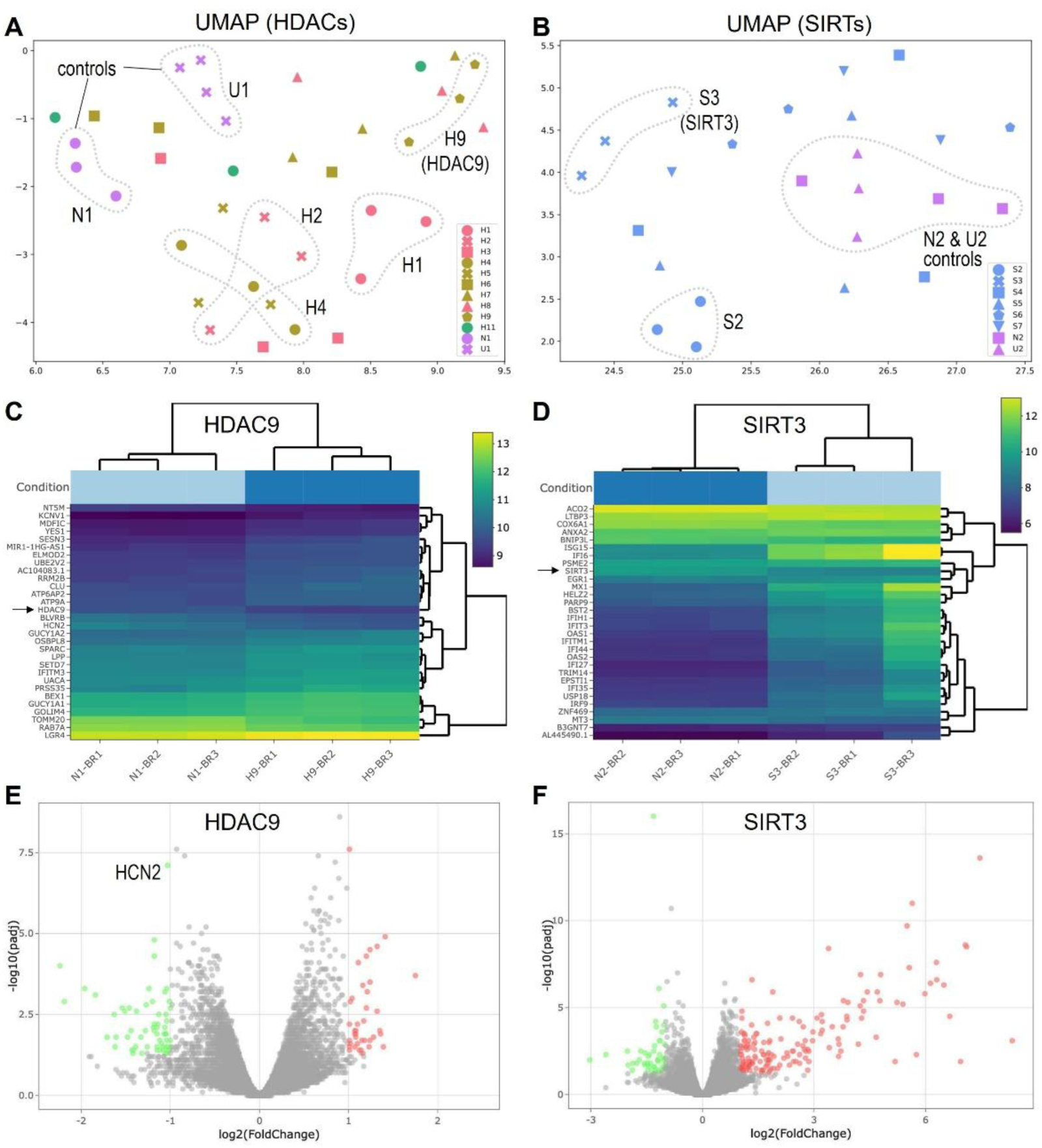
Whole-transcriptome (RNA sequencing) UMAP clustering of hiPSC-CMs treated by siRNAs against HDACs. **(A)** Results for targeting HDAC classes I, II, and IV **(B)** Results for targeting HDAC class III. UMAP embedding performed after univariate data imputation and scaling. Samples color-coded by HDAC class. N1 and N2 are negative controls treated by scramble siRNA on Plate 1 and Plate 2 respectively. U1 and U2 are untreated controls on Plate 1 and Plate 2 respectively. Dotted lines mark samples that clustered relatively tight and far away from the control groups. For U1 group, n=4; for all other treatment groups, n=3. Bi-clustering heatmaps display log2 transformed expression values, sorted by their adjusted p-value for top 30 differentially expressed genes between the HDAC9 knockdown group and corresponding negative control (**C**) and between the SIRT3 knockdown group and corresponding negative control (**D**). Volcano plots display global transcriptional change across HDAC9 knockdown group and negative control (**E**) and across SIRT3 knockdown group and negative control (**F**). Each dot represents one gene. Genes with adjusted p-value < 0.05 and log2 fold change > 1 are colored red and are considered up-regulated. Genes with adjusted p-value < 0.05 and log2 fold change < −1 are colored green and are considered downregulated in the knockdown group compared to the control.

Heatmaps for differential expression confirmed that targeted genes were among the top differentially expressed, demonstrated for *HDAC9* knockdown and *SIRT3* knockdown groups as examples (**Fig. 4C and 4D**). These heatmaps also identify other DEGs like *HCN2*, which was downregulated in the *HDAC9* knockdown samples (**Fig. 4C and 4E**). The volcano plots (**Figs. 4E and 4F**) allow further exploration of DEGs and demonstrate the overall effect of a particular knockdown. For example, *SIRT3* knockdown was associated more strongly with upregulation than downregulation of other genes, including many immunomodulatory factors, as suggested by earlier studies (53). Gene ontology results (**Fig. S2**) offer further lists of biological processes affected by *HDAC9* and *SIRT3* knockdown. Note that among the most affected processes were metabolism, membrane transport and excitability (for downregulation of *HDAC9* and *SIRT3*), and immune responses and viral entry (for downregulation of *SIRT3,* **Figs. 4D and 4F**).

### Regression modeling

Next, we deployed a method for linking *HDAC/SIRTs* to ion channel gene regulation through transcription factors as mediators (42). To do this, we structured three complementary partial least-squares (PLS) regression models: one in which *HDAC/SIRT* expression is the input and cardiac ion channel expression is the output (PLS1), a second, in which *HDAC/SIRT* expression is the input and *TF* expression is the output (PLS2), and a third, in which *TF* expression is the input and cardiac ion channel expression is the output. These models were built once using RNAseq data from the GTEx dataset and again, separately, using RNAseq data from our *HDAC/SIRT* knockdown hiPSC-CM. The suppression of each individual *HDAC/SIRT* gene in the hiPSC-CMs was aimed at creating a dynamic range to better inform relationships and downstream analysis.

### Modeling relationships between HDACs/SIRTs and ion channels

In comparing *HDAC/SIRT* genes to cardiac ion channel genes, correlation patterns emerge. For example, *HDAC/SIRT* and *HAT* genes are overall more positively correlated to ion channel genes in hiPSC-CM than in the adult LV (**Fig. 5A**). In both sample types, *HDAC7* and *HDAC10* show distinctly negative correlations with most ion channel genes. Certain “output” genes, like *KCNJ2, HCN2* in hiPSC-CM and *ATP1A1, HCN2* in adult LV, tend to be negatively correlated with “input” genes. Interestingly, *SIRT6, SIRT7* and *KAT2A* were strongly negatively correlated to ion channel genes in the adult LV but not in hiPSC-CMs.

**Figure 5.**
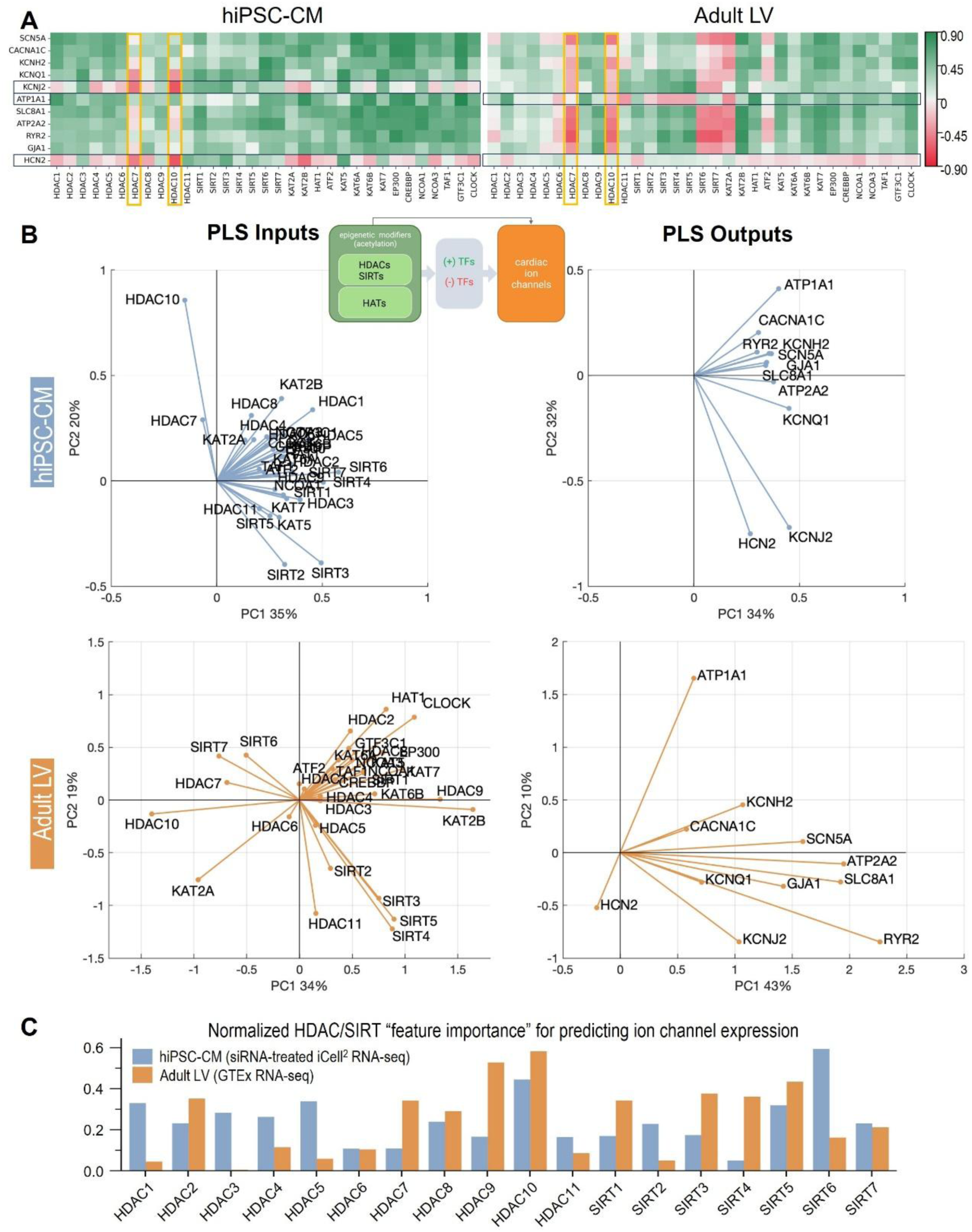
Analysis of relationships between HDAC/SIRT genes and ion channel genes in hiPSC-CM and adult LV bulk RNA-sequencing data. **A.** Correlation matrices where positive/negative correlations are colored green/red respectively. **B.** Results from PLS regression models are presented as biplots for the model inputs and outputs (HDACs/SIRTs and ion channels, PLS1) for hiPSC-CMs and for adult LV. Data for adult LV are modified from Pressler et al.(42). **C**. Average relative feature importance of HDAC/SIRT inputs in predicting ion channel outputs are displayed for hiPSC-CM and adult LV PLS models.

To further explore the relationships beyond correlations, the PLS models were constructed and their characteristics studied. We used biplots of the inputs and outputs to display PLS results in reduced dimensions, listing what fraction of the variance can be explained by each principle component. In all cases, the presented dimensions explained over 50% of the model results. Similarity of action or co-regulation of genes (i.e. simultaneous increase or simultaneous decrease) is represented in these biplots by a small angle between their projections; and the length of the lines indicates the strength of the gene’s effect in the model.

An important observation from PLS1 is that in both hiPSC-CM and adult LV, many of the key cardiac ion channel genes have small angles between them, suggesting co-regulation by the input (*HDAC/SIRT*) genes (**Fig. 5B**). This is true in the adult LV as well, where many input genes point in the same direction as the output genes, indicating a positive relationship between inputs and outputs. Within outputs, *ATP1A1*, *HCN2*, and *KCNJ2* are separated from other ion channel genes, which suggests being differently regulated than the others. A known co-regulation of genes encoding key depolarizing and hyperpolarizing currents *CACNA1C* and *KCNH2* (42, 43), believed to play a stabilizing, anti-arrhythmic role, is also seen in the adult LV and hiPSC-CMs. Similar co-regulation is seen for *ATP2A2* (SERCA2) and *SLC8A1* (Na/Ca exchanger) in both the adult LV and in the hiPSC-CMs.

In evaluating the general importance of each *HDAC/SIRT* gene in determining ion channel expression, we compared the “feature importance” parameter of the hiPSC-CM and adult LV PLS models (**Fig. 5C**). For hiPSC-CM, *HDACs 1-5, 9, 10,* and *SIRT6* expression had the largest impact on model performance. For the adult LV, *HDACs 9* and *10* seem more predominant. Note that higher values here do not necessarily indicate high absolute importance. Rather, they identify that a particular *HDAC*/*SIRT* is more important than the other *HDAC*s and *SIRT*s in driving the gene expression of cardiac ion channels.

### Modeling relationships between HDACs/SIRTs and transcription factors

Correlations between *HDACs/SIRTs* and *TF*s show interesting similarities among hiPSC-CM and the adult LV (**Fig. 6A**). For example, many *HAT*s are positively correlated with key cardiac TFs, especially *MEF2A*, *NFAT5*, and *NFKB1*. From the PLS2 analysis, both hiPSC-CM and adult LV showed clustering of *GATA4*, *GATA6*, and *MEF2A*, among others, suggesting similarity of action of those TFs (**Fig. 6B**)(54). *HDAC9* and *MEF2A* are co-aligned in both hiPSC-CM and adult LV, corroborating the well-studied interaction of *MEF2A* and *HDAC9* (30, 54). *HDAC9* is co-aligned with several critical TFs like *MEF2A*, *GATA4*, *NKX25*, and *YY1* (**Fig. 6B**).

**Figure 6.**
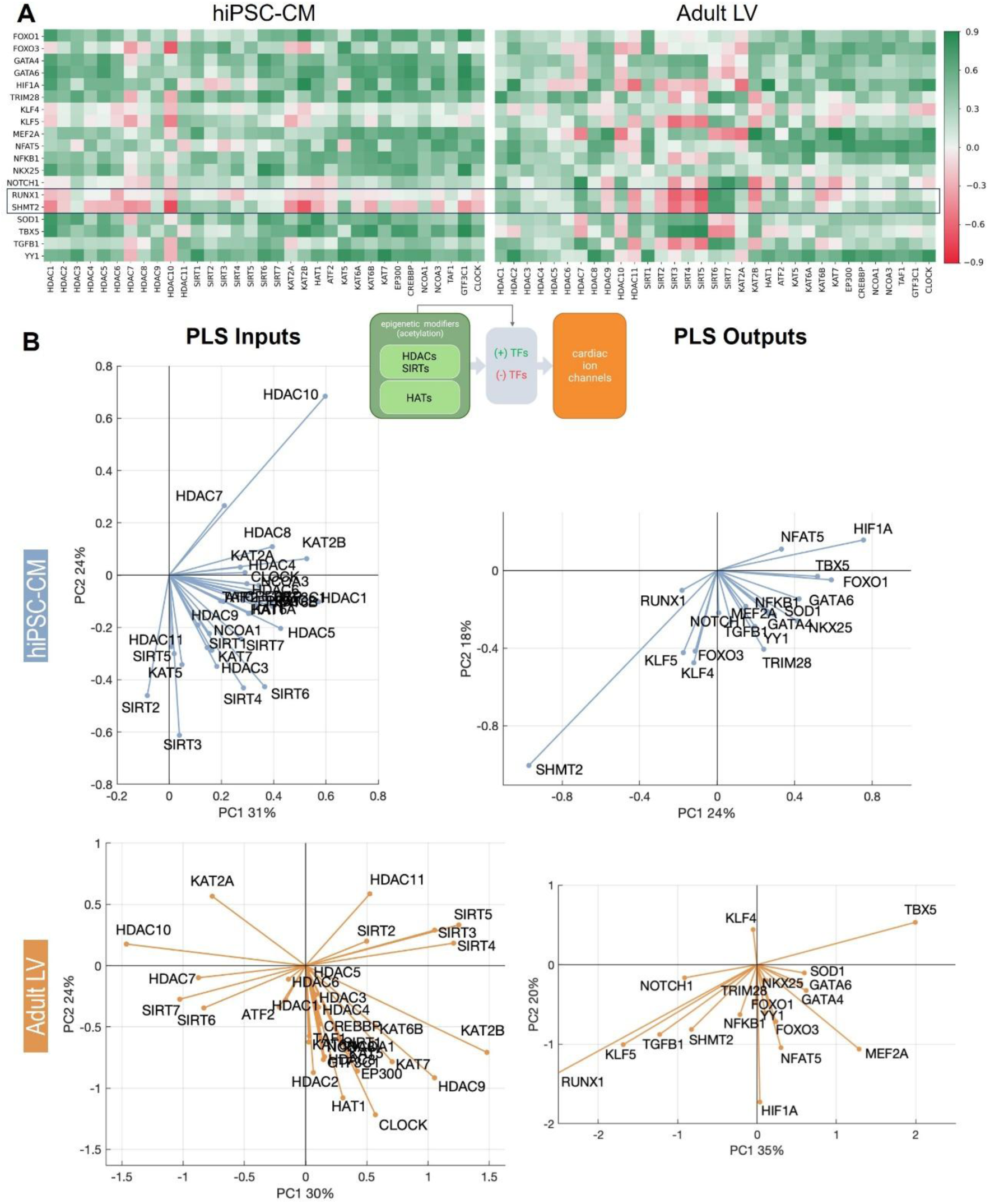
Analysis of relationships between HDAC/SIRT genes and TF genes in hiPSC-CM and adult LV bulk RNA-sequencing data. **A.** Correlation matrices show positive/negative correlations colored in green/red respectively. **B.** Partial least squares regression models are presented as biplots for the model inputs and outputs (HDACs/SIRTs and TFs, PLS2). Data for adult LV are modified from Pressler et al. (42).

### Modeling relationships between transcription factors and ion channels

We saw generally positive correlations between these gene sets in the hiPSC-CM, and both positive and negative correlations in the adult LV (**Fig. 7A**). A strikingly different role is seen for *TRIM28*, which has very high positive correlations with ion channels in hiPSC-CMs, but is mainly a negative regulator of ion channel expression in the LV. Similar differential roles are seen for *NOTCH1* and *TGFB1* and could be related to the distinct involvement of these TFs in early development vs. adult cardiac function. PLS3 shows strong alignment of many TFs and ion channels in hiPSC-CM (**Fig. 7B**) and suggests more complex relationships in the adult LV. *MEF2A* is an overall positive regulator of ion channel genes, as are *HIF1A* and *ATP1A1* in both sample types (**Fig. 7B**). Interestingly, PLS3, like PLS1, shows flanking by *ATP1A1*, *KCNJ2*, and *HCN2*.

**Figure 7.**
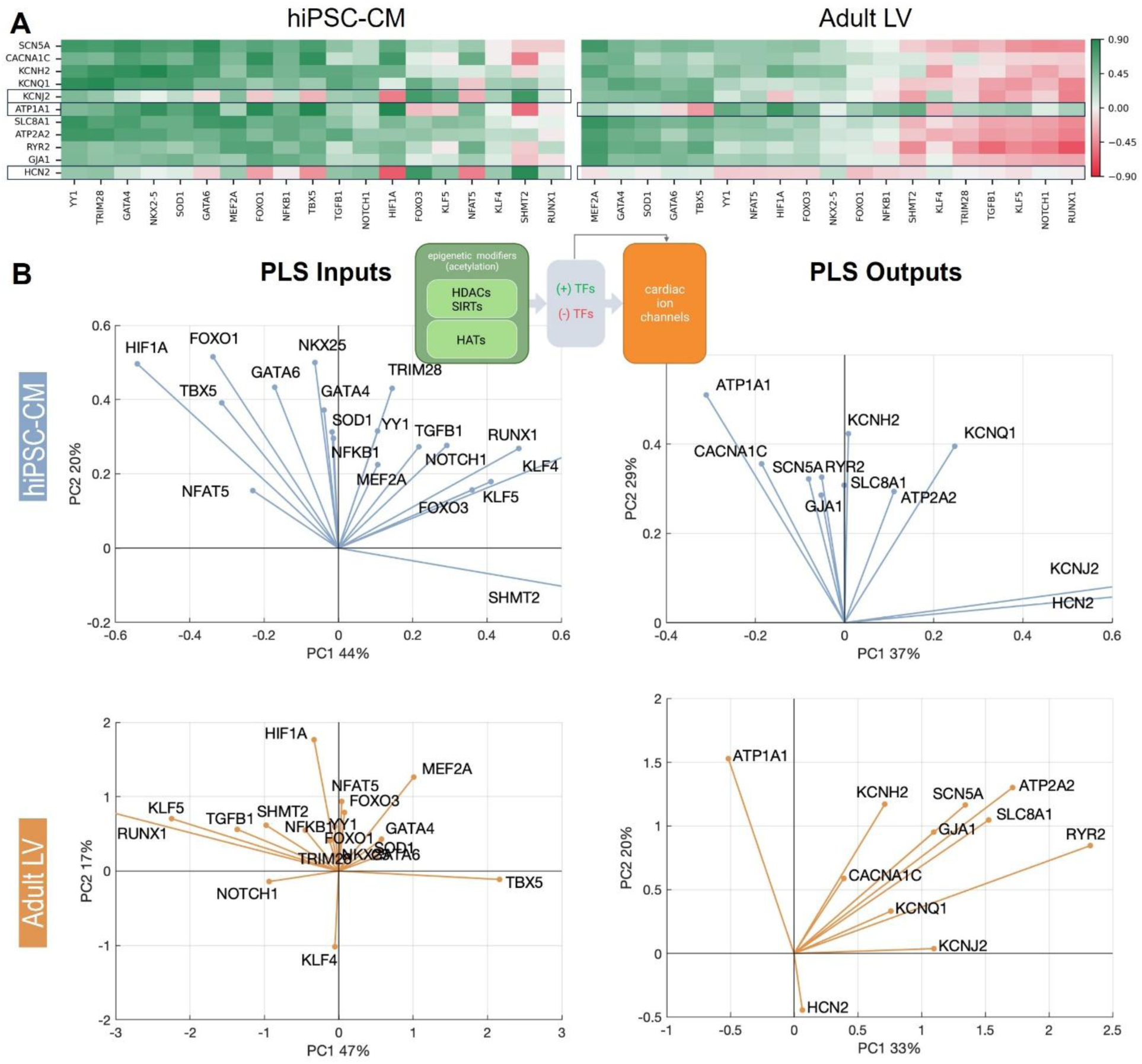
Analysis of relationships between TF genes and ion channel genes in hiPSC-CM and adult LV bulk RNA-sequencing data. **A.** Correlation matrices show positive/negative correlations colored in green/red respectively. **B.** Partial least squares regression models are presented as biplots for the model inputs and outputs (TFs and ion channels, PLS3). Data for adult LV are modified from Pressler et al.(42).

PLS3 also suggests some key differences between hiPSC-CM and adult LV. For example, similar to **Fig. 5B**, *KCNH2* and *CACNA1C*, which have opposite effects on action potential morphology, are closely aligned in the adult LV (**Fig. 7B**) but less so in the iPSC-CM. This suggest stronger co-regulation by *HDACs/SIRTs* (**Fig. 5B**) and by cardiac TFs (**Fig 7B**) in the adult LV than in hiPSC-CM. Likewise, *KCNJ2* and *HCN2* are opposite regulators of pacemaking and are co-aligned in hiPSC-CM but not in adult LV. Several TFs that show strong influence in hiPSC-CM (large magnitude) appear less impactful in the adult LV, such as *TRIM28*, *FOXO1*, *NKX25*, and *GATA6*.

### Sankey plots illustrate detailed links between inputs and outputs

An alternative approach for analyzing the PLS regression models allows a display of all three models in conjunction for both the hiPSC-CM (**Fig. 8A**) and adult LV (**Fig. 8B**). Predicted positive (green) and negative (red) regulators of ion channel expression can be easily viewed here, with the strength of relationships encoded by the line thickness. Major positive regulators like *GATA4*, *GATA6*, *HIF1A*, and *TBX5* are seen in hiPSC-CM and adult LV. *FOXO1*, *NFAT5*, and *SHMT2* seem to be stronger regulators in hiPSC-CM than in the adult LV. *TRIM28* seems to be a positive regulator of ion channel genes in the hiPSC-CMs but a negative regulator in the adult LV. *MEF2A* (a strong positive regulator) and *RUNX1* (a strong negative regulator) are shown to be more active in the adult LV than in hiPSC-CM. Overall, TFs in hiPSC-CM tend to be predominantly positive regulators of ion channel expression compared to TFs in the adult LV.

**Figure 8.**
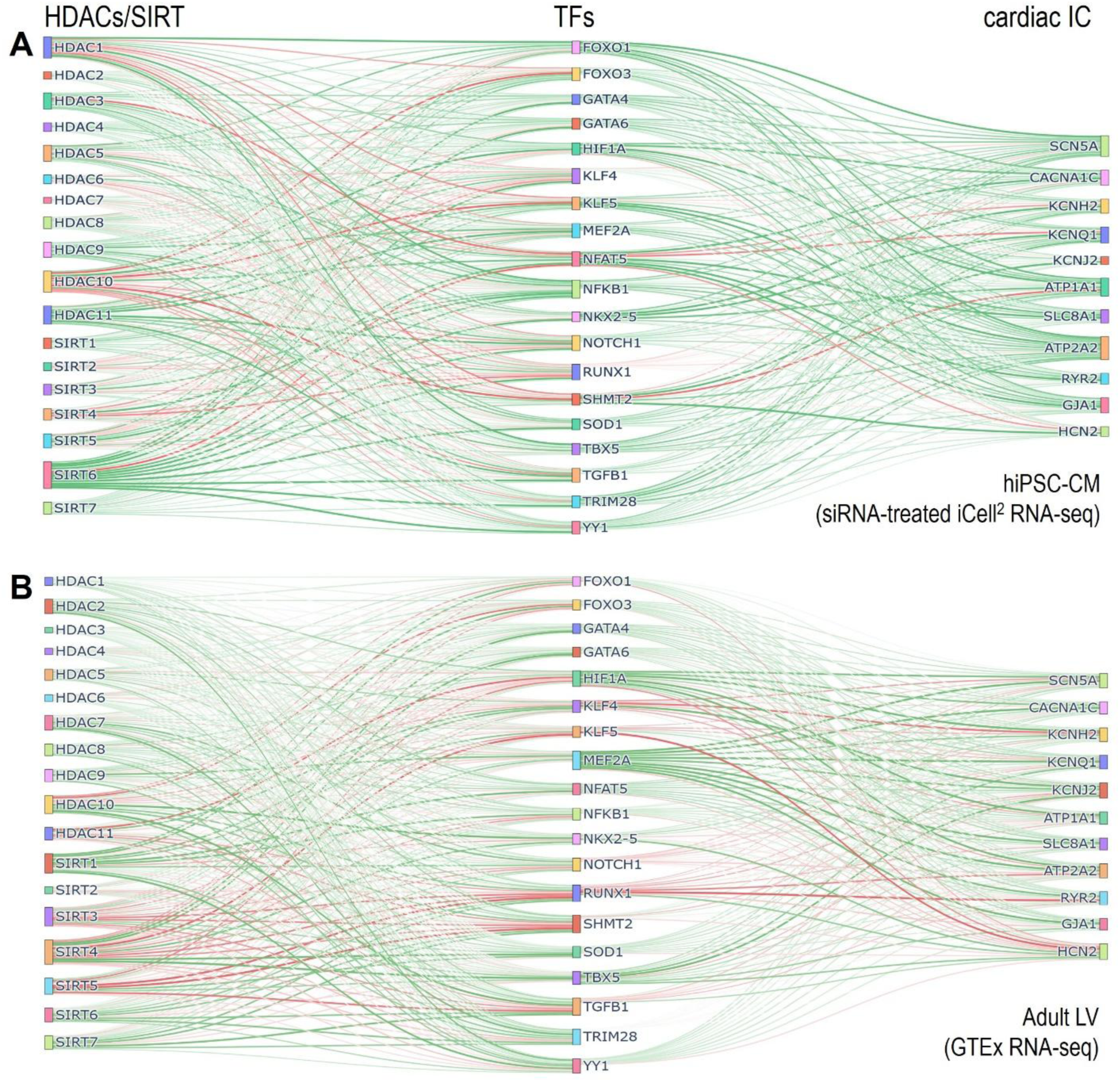
Links between HDACs/SIRTS, TFs and cardiac ion channels. Integrated (Sankey) visualizations depict links between HDACs/SIRTs (left), TFs (middle), and ion channels (right) in hiPSC-CM (**A**) and adult LV (**B**). Color and weight of lines defined by sign and magnitude of the PLS β-matrix coefficients.

### Spontaneous beat rate after *HDAC*/*SIRT* knockdown

Given the large number of complex and overlapping predictions made by the PLS models, we attempted preliminary explorations of HDAC and SIRT impact in cardiomyocyte electrophysiological function. Over many successive cell thaws, spontaneous beat rate in hiPSC-CM was measured for all *HDAC* and *SIRT* knockdowns (**Fig. 9A**, stratified by class in **Fig. 9B**). Most apparent was a general increase in spontaneous beat rate after *HDAC* knockdowns and a general decrease in spontaneous beat rate after *SIRT* knockdowns, **Fig. 9A**. Inhibition of HDAC2 and HDAC10 had the strongest effect – increasing excitability and spontaneous rate. There were exceptions, such as *HDAC7* and *HDAC9* (members of HDAC class IIa), which, when knocked down by siRNA, decreased beat rate and excitability, similar to the SIRTs. Interestingly, knockdown of *SIRT2* resulted in a powerful reduction of beat rate that was even stronger than the effect of propranolol, a negative rate control. Indeed, many *SIRT2* knockdown samples were completely quiescent 48hrs after siRNA transfection. This agrees with the co-alignment of *SIRT2* and the pacemaking gene *HCN2*. Detailed statistics on the effect of *HDAC/SIRT* inhibition are shown in **Table S2**.

**Figure 9.**
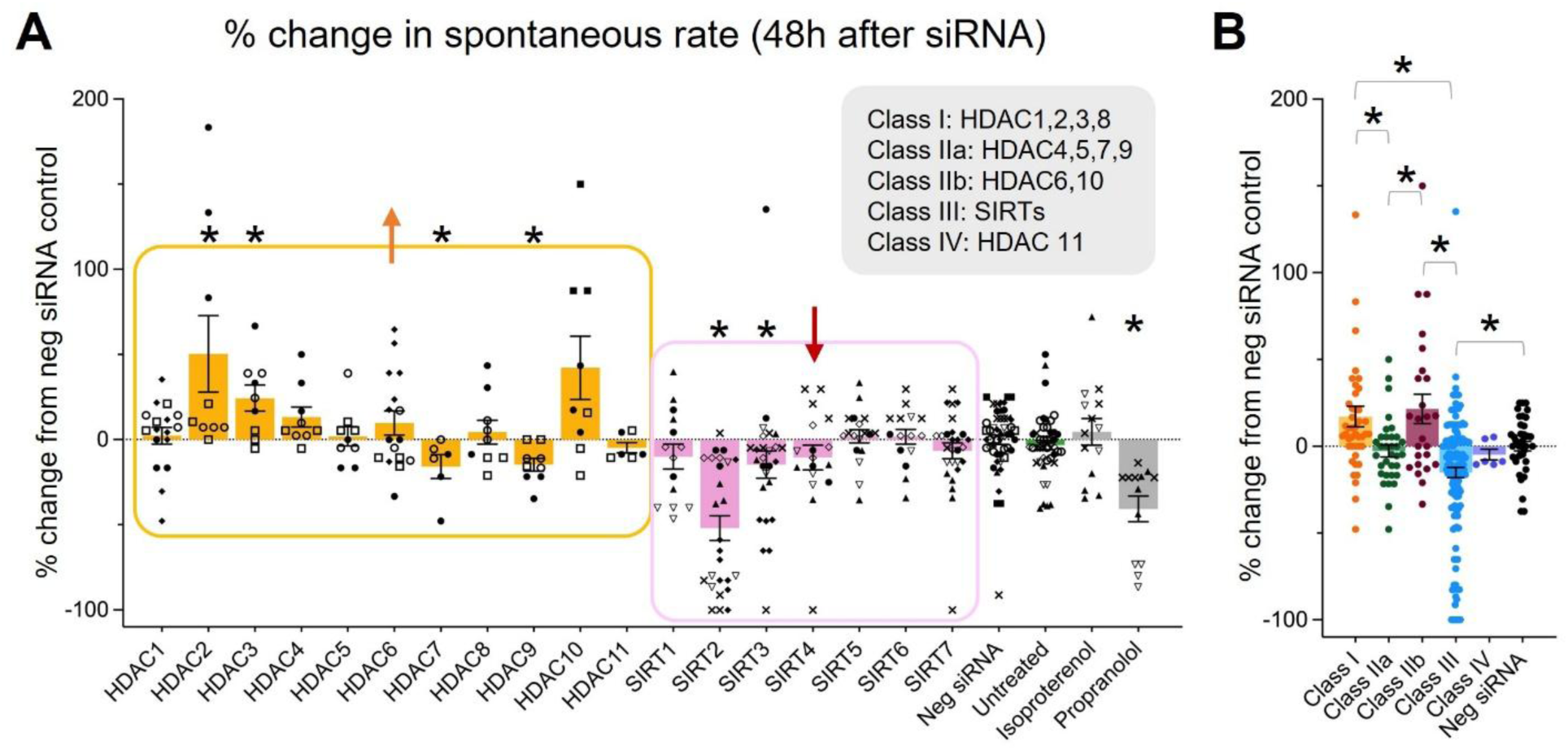
Modulation of excitability and spontaneous rates of hiPSC-CMs by suppression of HDACs and SIRTs. **A**. Spontaneous beating rates of hiPSC-CM treated for 48h with siRNAs against individual HDACs and SIRTs, displayed as percent change from the negative siRNA control. Isoproterenol and propranolol were included as positive and negative rate controls respectively, although after 48 h isoproterenol’s chronotropic action is almost completely absent. All error bars represent mean and standard deviation. Different symbols represent experimental repeats across ten separate cell thaws. In general, suppressing HDACs leads to overall increase in beat rate and excitability while suppressing SIRTs leads to overall decrease in beat rate/excitability. **B**. The same data grouped by class of HDACs (I, IIa, IIb, III and IV). Statistical significance (p < 0.05) in difference from negative siRNA (panel A) or between groups (panel B), found by non-parametric Kruskal-Wallis test, is indicated by an (*). n=6-143 per group.

## DISCUSSION

We present a direct comparison of deep sequenced transcriptomics data from the adult human heart and from iCell^2^, a popular commercial hiPSC-CM cell line, optimized for ventricular phenotype and widely used in pharma and academia. This study used siRNA gene modulation of epigenetic modifiers in conjunction with partial least square models to investigate transcriptomic relationships between *HDAC/SIRT*s, cardiac TFs, and ion channel genes. Our motivation to focus on these genes was the interest in using the hiPSC-CMs as a preclinical experimental platform for cardiotoxicity studies for new compounds including HDACi, where cardiac arrhythmia problems are of prime concern.

We found strikingly similar patterns of expression for key cardiac ion channel genes in the hiPSC-CM and the adult human LV. While transcriptional ion channel similarities have been reported between iCell^2^ and isolated human ventricular myocytes (21), our results are quite surprising considering the adult LV tissue contains a roughly half and half mixture of myocytes and non-myocytes (55). One plausible explanation is that ventricular myocytes are likely the main contributors to the detected ion channel transcripts. Indeed, previously published single-nuclei sequencing of heart tissue shows that ion channel transcription in non-myocytes, e.g. fibroblasts, is minimal compared to cardiomyocytes (52).

Cardiac TFs also exhibited high similarity of expression between the hiPSC-CMs and the adult LV, possibly because of cardiomyocyte-specific expression. The expression patterns of *HDACs/SIRTs* and *HATs* in the hiPSC-CM and the adult LV were similar but to a lesser degree than the ion channels and the TF genes which, considering the dynamic nature of these epigenetic players, is noteworthy.

Perturbing *HDACs/SIRTs* expression in the hiPSC-CMs allowed a wider dynamic range of the dataset and we used our previously developed PLS regression strategy to compare key relationships with electrophysiology-relevant genes (42). The PLS models suggest substantial co-regulation of the cardiac ion channels both in the adult LV and in the hiPSC-CMs. In other words, ion channel genes are upregulated or downregulated in tandem by key cardiac TFs and by *HDACs/SIRTs*. This finding is in line with recent studies reporting highly coordinated regulation of ventricular ion channel expression (43) and co-occupancy of genomic regions by multiple cardiac TFs, operating synchronously (54) to achieve stability amidst dynamic epigenetic modulations (42, 43). Furthermore, the analysis suggests unique regulatory patterns for ion channels that are critical for maintenance of the resting membrane potential, e.g. *ATP1A1, KCNJ2,* and *HCN2.*This may be a manifestation of compensatory mechanisms to maintain electromechanical homeostasis (42), both in the hiPSC-CM and adult LV.

The PLS analyses also highlight positive and negative regulators of cardiac ion channel expression. The known positive regulation of cardiac TFs like *GATA4, GATA6, MEF2A*, and *HIF1A* (54, 56) is captured in our analysis for both hiPSC-CM and adult LV. Our reported co-alignment of *HDAC4*,*5*, and *9* with *MEF2A* in hiPSC-CM and the adult LV is congruent with the fact that class IIa HDACs (except *HDAC7*) have a *MEF2*-binding domain (57, 58) and that *MEF2A* is a strong positive regulator of ion channel gene expression, hence why class IIa HDACs are considered protective in the heart (59).

*SIRTs* are a special HDAC type (class III) with a metabolism-sensing domain (NAD+). They are also perceived as cardioprotective and as positive regulators of cardiac ion channels (53, 60–64). This is most obvious in the strong co-alignment we saw of mitochondrial sirtuins *SIRT3-5* and cardiac ion channels, especially in the adult LV. However, our results suggest differential roles for the nuclear sirtuins *SIRT6, 7* in modulating cardiac ion channels, which remains to be explained.

The PLS models suggest that *FOXO1* is more active in hiPSC-CM than in the adult LV, which is consistent with reports that the *FOXO* family, regulators of proliferation, are not active in the adult LV, as it has transitioned to hypertrophy (65). Distinct roles were also seen for *TRIM28* (also known as *KAP1*) – as a predominantly negative regulator of ion channel genes in the adult LV, yet a positive regulator of ion channel genes in the hiPSC-CMs. Recent work from our group indicates that this may be important for designing activator and suppressor domains for CRISPRa/i (activation/interference). *TRIM28* plays a key role in partnering with KRAB-ZFN (Krueppel-associated box - zinc-finger) domains used in CRISPRi (66). Our recent report found that Zim3 (a potent KRAB-ZFN domain for CRISPRi (67)) paradoxically acts as an activator (instead of inhibitor) of ion channel genes in hiPSC-CMs (68).

Generally, these marked differences could be due to the sample sources – hiPSC-CM are stem cell derived while the adult LV tissue from the GTEx dataset are samples retrieved postmortem. Transcriptional differences could also be explained by the sample composition, comparing pure cardiomyocytes to multi-cell-type tissue samples, particularly for genes without cardiomyocyte-specific expression. Unfortunately, read depth of single cell (scRNAseq) or single nucleus (snRNAseq) sequencing, in most currently available transcriptomic datasets, is insufficient for detailed exploration of the gene sets considered here. Our bulk RNAseq was performed using densely-grown hiPSC-CM syncytia, which we deem essential for transcriptomics results, as previous work has shown that cell density is a key factor of ion channel expression (46) and broader transcriptomic outcomes, with adult-like features missing when cardiomyocytes are grown sparsely (20). This should be considered in sample processing for future single-cell/single-nucleus studies, especially for genes with fast dynamics of expression. Importantly, other researchers have found that for cardiomyocyte-specific genes, 3D vs. 2D growth of pure iPSC-CMs in dense monolayers did not drastically change the transcriptomics results (22); in contrast, 3D vs. 2D growth led to dramatic transcriptomic changes for non-myocytes, e.g. endothelial cells, fibroblasts or immune cells (22).

There are several limitations of the presented approach. PLS modeling is good at generating predictive input-output relationships because its strength is to uncover/maximize covariances (69). However, causality in such relationships may still be difficult to prove because the structure of our assumed model is simplified, without consideration of complex feedback loops. Gene regulatory networks capitalizing on prior knowledge of key relationships may be an alternative approach to consider in future. Another limitation of focusing exclusively on transcriptomics data is that RNA transcript levels do not always correlate with protein levels and, ultimately, with functional performance. Therefore, the functional relevance of novel relationships suggested by the PLS model must be confirmed experimentally.

Here, we optically quantified of the effects of *HDAC/SIRT* suppression on spontaneous beat rate. Suppression of *HDACs,* in general, increased excitability while suppression of *SIRTs* generally decreased excitability. Our results suggest a possible mechanism for these effects: the negative regulation of HCN channels by *HDAC* classes I, II, and IV implies that *HDAC* inhibition may lead to increased pacemaking, while positive regulation of *HCN2* by sirtuins implies that *SIRT* inhibition may suppress pacemaking, **Fig. 5**. From our analysis, *SHMT2* and *NFAT5* (positive and negative *HCN2* modulators, respectively) may be implicated for these effects in hiPSC-CMs, **Figs. 7** and **8A**. Perhaps a better known relationship is *SIRT1*’s positive regulation of *SCN5A* and deacetylation-mediated trafficking of sodium ion channels (70), which might explain in part the overall reduction of excitability when *SIRT*s are suppressed. The distinct effects of HDACs on excitability, individually and stratified by class, are informative for HDACi design. Comprehensive functional testing beyond heart rate is needed to confirm transcriptomics-based predictions on the action of HDACis on cardiac electrophysiology.

Future work would benefit from an interference CRISPR approach for gene modulation, followed by all-optical electrophysiology, as we have previously shown for several key ion channel genes (9). CRISPRa/i would allow investigation of both loss-of-function and gain-of-function perturbations (8, 9). Such elaborate datasets, along with detailed functional characterization, may better capture the complex interconnectedness of the studied molecular players. We view the PLS models as an exploratory tool to identify regulatory links, with specific mechanisms being later verified by chromatin immunoprecipitation sequencing (ChIP-seq) or assay for transposase-accessible chromatin using sequencing (ATAC-seq). Indeed, the combination of predictive modeling, all-optical functional measurements, and ChIP-seq or ATAC-seq would be highly informative in dissecting these complex relationships.

In summary, this study demonstrates similarities in gene expression patterns and in transcriptional regulation of ion channel genes by *HDACs/SIRTs* and key cardiac TFs between an optimized commercial hiPSC-CM cell line (iCell^2^) and adult human heart tissue from the GTEx dataset. We corroborated multiple relationships known from animal studies, and future in-depth functional testing will help further validate human iPSC-CM as a translational, scalable platform for cardiac epigenetic studies and for development of new cardiac therapeutics.

## METHODS

### Cell culture and sample preparation

Human iPSC-derived cardiomyocytes (iCell Cardiomyocytes^2^ from Fujifilm Cellular Dynamics International) were thawed according to manufacturer’s protocol and plated in fibronectin-coated glass-bottom plates. Samples cultured for RNAseq were plated in 48-well format at 150,000 cells/well. Samples cultured for quantification of spontaneous beat rate were plated in 96-well format at 50,000 cells/well. For all cardiomyocyte culture, maintenance media was refreshed every two days.

### Gene knockdown by siRNA transfection

siRNAs against *HDAC* and *SIRT* genes and a scrambled negative control (MISSION® esiRNA, Millipore Sigma or Silencer® Select, ThermoFisher) were transfected according to manufacturer’s protocol on cell culture day 5. Details on the siRNAs used are provided in **Table S1**. siRNA constructs and transfection reagent were applied at 150ng/well and 3μL/well, respectively, for 48-well format and at 50ng/well and 1μL/well, respectively, for 96-well format. Transfection media was replaced with normal maintenance media on cell culture day 7, after 48 h of treatment. Each culture plate included untreated (non-transfected) controls and media was refreshed every two days. Samples were collected for RNAseq on culture day 7.

### Sample preparation and RNA-sequencing

Samples for RNAseq (total n=61) were washed with cold RNAase-free PBS. To each sample, 50 µl Trizol was applied, cells were lysed, homogenized and collected in cryo-vials to be flash-frozen in liquid nitrogen prior to shipping. The target RNA per sample was 400 ng. Samples were sent to GENEWIZ/Azenta, South Plainfield, NJ, and sequenced using Illumina pe150 platform to a target depth of 50 million reads and 150-bp paired-end fragments.

### Data processing and analysis

RNAseq data for adult human hearts (left ventricle, LV) were accessed from the GTEx dataset (23) and preprocessed as described by Pressler et al.(42). Briefly, expression levels in transcripts per million (TPM) in heart left ventricle samples were extracted from the GTEx v.8 dataset. These RNAseq libraries were generated by Illumina TruSeq protocol (non-strand specific, polyA-based). Median depth was 78 million 76-bp paired-end reads. Human reference genome Homo_sapiens.GRCh38.79.gtf was used. TPM values were filtered by *GAPDH* (including only samples with *GAPDH* TPM between 500-2500), normalized by geometric mean and log-ratio representation (71, 72). To match the hiPSC-CM cell line, which is derived from a female donor, we analyzed only data from female samples (n=84, the same set as in Pressler et al.), which exhibited natural variablility due to age (21 to 70 y.o.) and other health status and phenotypic characteristics. These GTEx data subset were processed in an identical way as the hiPSC-CM RNAseq data to allow direct comparison.

### Quantifying similarity of gene expression patterns

In **Figs. 2-3**, we display patterns of gene expression for the genes of interest in hiPSC-CMs and adult LV. Radial plots of the mean values from the corresponding bar charts help visualize similarity. Furthermore, we calculated a metric for each gene set to quantify how different the gene expression patterns were in hiPSC-CM vs. the adult human LV. For each gene of interest, the difference between expression values in hiPSC-CM and LV were normalized to the magnitude of the expression values (Eq. 1). High k values correspond to high difference and equal expression values would result in a k value of zero. Difference scores K_difference_ are the median of k values for each gene set.

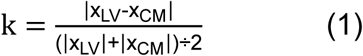

### Clustering

RNAseq dataset from siRNA-treated hiPSC-CM underwent univariate imputation after 634 genes with blank values from all samples were removed. Expression data were then scaled to zero mean and unit variance and embedded to UMAP using n_neighbors = 10 and all other parameters set to default. Because the samples depicted in the two UMAPs were comprised of two separate batches, batch correction was not necessary.

### Correlation analysis

In both the hiPSC-CM and adult LV RNAseq datasets, Pearson’s correlations were calculated between *HDACs/HATs* and ion channels, *HDACs/HATs* and transcription factors, and transcription factors and ion channels.

### Partial least squares regression models

Three PLS models each were built for the hiPSC-CM and adult LV data: PLS1, considering *HDACs/HATs* as inputs and ion channels as outputs; PLS2, considering *HDACs/HATs* as inputs and transcription factors as outputs; and PLS3, considering transcription factors as inputs and ion channels as outputs. As described in Pressler et al.(42), these models and their predictions were calculated in Python using PLSRegression from the sklearn library and the number of components was determined by leave-one-out cross validation and predicted residual error sum of squares. To explain the most variance while also mitigating overfitting, 4 components were selected for these PLS models.

### Visualization by biplots

PLS model inputs and outputs are presented as “biplots”, where the PLS loadings are projected onto lower dimensional space of the first two latent variables of the 4-latent-variable model. The axes indicate the percent variance explained by these two latent variables. The magnitude of each line in the biplots denotes the importance of that variable in the projection and the alignment or proximity of angle between variables signifies similarity of action. Importantly, these biplots represent the average results from Monte Carlo Cross-Validation, where 1,000 runs were performed with random train-test splitting (with 10% holdout). The resulting loadings were averaged across the runs and projected in **Figs. 5-7**.

### Feature importance model inspection

The importance of *HDAC/SIRT* model inputs in potentially influencing ion channel outputs was quantified by the permutation importance of the PLS regression inputs. Permutation importance is a metric of how much each model “feature” (i.e. input) contributes to the predictive performance of the model. In the sklearn library, this is calculated by permuting feature values (by random amounts) of a single feature and measuring the effect on the model score. PLS models were constructed with *HDAC*/*SIRT* as inputs and a single ion channel gene as the model output. This was repeated for each of the ion channel genes discussed here. The permutation importance for individual ion channels was extracted and normalized to 1. These relative values were then averaged across the eleven ion channel outputs so that the feature importance displayed for each *HDAC*/*SIRT* input represents a generalized importance in predicting ion channel expression.

### Visualization by Sankey plots

Sankey plots depict new PLS models trained by the hiPSC-CM and adult LV RNAseq data, separately and with zero holdout, using the 4 components previously selected. To avoid overshadowing of small-scale relationships between HDACs/SIRTs, TFs, and ion channels, for each of the performed PLS simulations, the beta matrix coefficients were scaled to be between −1 and 1, with no center shift, using MaxAbsScaler from the sklearn library. These scaled beta matrices informed the plots: the sign of matrix coefficients determined line color and the coefficient magnitudes determined line weight, which signify the strength and direction of the respective relationships.

### Quantification of spontaneous beat rate with individual HDAC/SIRT inhibition

After 48-hour treatment with siRNA or controls, hiPSC-CM monolayers in 96-well format were labeled with voltage sensitive dye BeRST1 (3, 73) and imaged on an in-house developed OptoDyCE-plate macroscope system (6, 9). This allowed simultaneous data capture from all experimental groups. Beat frequencies were normalized against negative siRNA control within the same plate so that data from multiple experiments could be combined. Isoproterenol (or isoprenaline) at 10 µM and propranolol (10 µM) were included as positive and negative rate controls, respectively, and were applied 24 hrs before imaging and in some cases again immediately before imaging.

### Statistical Analysis

Statistical analysis was performed in GraphPad Prism (v.10.1.1, GraphPad Software, Boston, MA). Differences between spontaneous rates after siRNA treatments to inhibit different *HDACs/SIRTs* were deemed significant at p<0.05 assessed by a non-parametric Kruskal-Wallis test and pairwise comparisons.

## Author contributions

EE and MRP designed the study. MRP conducted experiments, analyzed data, and organized all figures. MPP and MRP developed and applied computational analysis tools. AH pre-processed the RNAseq data and helped access and process the GTEx data. EE supervised the study and provided resources. MRP and EE wrote the manuscript with input from AH and MPP.

## Acknowledgements

This work was supported in part by a grant from the National Science Foundation EFRI 1830941 (EE), a fellowship from the National Institutes of Health 1F31HL168800-01 (MRP) and a grant from the Human Frontiers Science Program RGP010/2024 (EE).

## Declaration of interests

The authors declare no conflicts of interest, financial or otherwise.

## Data and materials availability

The raw data and the TPMs after alignment from the RNAseq in the hiPSC-CMs have been deposited at the National Center for Biotechnology Information Gene Expression Omnibus database under accession GSE309350. The RNAseq dataset from the adult human heart is from the GTEx repository: https://gtexportal.org/home/dataset. Custom software for analysis is referenced; the Python and MATLAB scripts are made available through github: https://github.com/pressm/PLSGTEx.

## Notes

### Competing Interest Statement

The authors have declared no competing interest.

